# Morphological details contribute to neuronal response variability within the same cell type

**DOI:** 10.1101/2025.09.15.676219

**Authors:** Kevin Sandbote, Ihor Arkhypchuk, Jutta Kretzberg

## Abstract

A large body of literature offers an explanation on how electrical diversity and variable branching patterns of neurons contribute to degeneracy, enabling multiple solutions for a characteristic neuronal response. For neurons with the same branching pattern it is yet unclear, how finer morphological details, such as diameter and length of dendritic branches, contribute to response variability. We address this question using a model database approach with spatially extended, conductance-based compartmental models to study variability of response features, such as resting membrane potential, input resistance, spike count, first spike latency, spike height, and spike width. Using 15 reconstructed morphologies of leech touch cells with fixed branching patterns, we identified several thousands of parameter sets that reproduced the experimentally measured response features in all the tested morphologies. Even when the electrical parameters were kept equal across reconstructed morphologies, variability in response features arose from the morphological details. This could not be explained by well-known dependencies on the total membrane area and input resistance. Systematically varying the spatial distribution of ion channels revealed that spike response features are influenced by the location of spike initiation zones with higher conductance density. Nevertheless, biologically plausible responses can arise from different locations of spike initiation zones and even homogeneous distribution of ion channels. Furthermore, comparing the simulated spike responses from two morphological subtypes of leech touch cells revealed that the previously published systematic differences cannot be explained by the morphological differences alone. A larger total conductance of voltage-gated ion channels was required to reproduce the experimental finding of an increased spike count and a larger spike amplitude in a larger morphological subtype. In conclusion, electrical properties, morphological details, and ion channel distribution across the membrane all interact in their contribution to the functionality and response variability of neurons of the same cell type.

## 1 INTRODUCTION

The brain is often compared to a computer, suggesting that its components exhibit algorithmic functionality and deterministic behavior (Brette, 2022). However, the idea that brains are built of many identical neurons is a fallacy, since cell-to-cell variability in neuronal responses is observed in nervous systems across species, e.g. in hippocampal neurons in rats (Rathour and Narayanan, 2019), the visual system in drosophila (Linneweber et al., 2020), the stomatogastric nervous system of lobsters (Bucher et al., 2005) and the nervous system of leeches (Calabrese et al., 2011; Norris et al., 2011; Roffman et al., 2012; Wenning et al., 2018; Scherer et al., 2022). In addition to this functional response variability, morphological variability in dendritic and axonal structure even within a defined cell type was reported by many studies (Goodman, 1978; Peng et al., 2021; Moubarak et al., 2022; Meiser et al., 2023). The multiple sources of neuronal variability do not necessarily impair neuronal functionality, because functional network behavior can result from various combinations of neuronal configurations as was shown in experimental and modeling studies (Prinz et al., 2004; Achard and De Schutter, 2006; Alonso and Marder, 2019). The concept of degeneracy, in which distinct structural configurations lead to similar functional outcomes, has emerged as a key principle exemplifying how biological systems maintain robust performance despite underlying variability (Edelman and Gally, 2001). In the context of neural circuits, degeneracy enables flexible computation and resilience to perturbations (Goaillard and Marder, 2021a; Schneider et al., 2023).

Computational modeling has long been a cornerstone of neuroscience research. Modeling approaches, such as the Hodgkin-Huxley formalism (Hodgkin and Huxley, 1952; Almog and Korngreen, 2016), often aimed to identify a single set of “optimal” parameters that reproduced a “typical” experimental observation. While powerful, such models do not always capture the broad range of observed biological variability. Increasingly, studies adopt a database modeling approach, in which large databases of conductance-based models were systematically examined to demonstrate that diverse combinations of ionic and synaptic properties can generate functionally similar network dynamics and neuronal outputs (Prinz et al., 2003, 2004; Gü nay et al., 2008; Marder and Taylor, 2011; Jedlicka et al., 2022). These findings led to the understanding of degeneracy and variability, even within cells of the same type. The excitability of an individual neuron is known to depend on its input resistance, which correlates negatively with the total membrane surface area (Kernell, 1966; Kernell and Zwaagstra, 1981) and is also affected by various types of ion channels (Ceballos et al., 2016, 2017). Modern computational studies have focused primarily on variability between different cell types and variability in dendritic branching patterns (Mainen and Sejnowski, 1996; Eyal et al., 2014; Kole and Brette, 2018; Goaillard et al., 2020; Moubarak et al., 2022). However, the role of morphological details beyond cell size and branching patterns in shaping variability in cell-to-cell responses of a specific cell type remains to be explored.

The leech nervous system provides a very accessible platform to investigate this question. It is traditionally considered stereotyped and consists of segmentally repeated ganglia with well-characterized and individually identifiable neurons (Kristan et al., 2005). The mechanosensory touch cell (T cell) encodes light tactile stimulation of the leech skin with its spike count and first spike latency (Pirschel and Kretzberg, 2016). While the same individual T cell responds very precisely to repeated skin stimulation, the responses vary widely between neurons of this cell type (Scherer et al., 2022, 2023). Early studies on leech mechanosensory neurons revealed, that T cells exhibit two distinct morphological subtypes, innervating different regions of the skin: T cells with one root process innervating the skin (1RP) and T cells with two root processes innervating the skin (2RP) (Nicholls and Baylor, 1968). More recent studies find that these morphological subtypes contribute to the cell-to-cell variability with systematically different response features (Meiser et al., 2023). Meiser et al. speculated that the systematically higher spike counts and larger spike amplitudes observed in 2RP cells are likely caused by a larger area of active membrane compared to 1RP cells (Meiser et al., 2023). This hypothesis cannot be tested experimentally because (to our knowledge) there are no established antibody staining methods for leech neurons capable of marking ion-channels or cell structures associated with spike initiation zones (SIZ), as was conducted previously in other species (Meeks and Mennerick, 2007; Guo et al., 2017). Therefore, the distribution of voltage-dependent ion channels across the membrane of T cells can only be inferred indirectly from experimental and modeling studies. Previous studies suggested that a T cell has at least two SIZs, one (or more) in their distal processes innervating the skin and one (or more) SIZ proximal to its soma within the ganglion (Burgin and Szczupak, 2003; Kretzberg et al., 2007). Furthermore, a recently published single-compartment model of the T cell was able to reproduce typical response features, but systematically failed to reproduce the experimentally observed first spike latency, suggesting that spikes are not generated in the soma (Meiser et al., 2019).

Hence, in this study we use a multi-compartmental modeling approach that incorporates spatial structure and ion channel distribution, to analyze their impact on spike generation and to explain systematic differences in response features between the two morphological subtypes. Combining reconstructed morphologies, electrophysiological recordings, and conductance-based modeling, we assess how fine-scale morphological differences affect a cell’s response to electrical stimulation. Furthermore, our study sheds light on systematic differences in the response patterns of morphological subtypes of the same cell type. By varying electrical parameters and analyzing responses from a population of reconstructed morphologies, we isolate morphological variability from electrical factors, and aim to clarify its specific contribution to functional variability.

## 2 RESULTS

Leech T cells typically respond with short bursts of action potentials when their soma is electrically stimulated. However, the number, timing and shape of the action potentials vary between individual T cells. In order to investigate the influence of detailed morphological diversity on neuronal response variability, we generated and analyzed a large database of parameter sets that produced biologically plausible responses when applied to the reconstructed morphology of a leech T cell, rather than minimizing specific error values for the compartmental model. We defined a parameter set as *valid* and its model output as *biologically plausible*, if all analyzed response feature values were within the experimentally observed range based on 60 intracellular recordings of T cells (Meiser et al., 2023). The resting membrane potential and the input resistance were considered as subthreshold response features. The spike features were determined in response to a 500 ms long current injection of +1.5 nA into the soma. They comprised the spike count, the first spike latency, the spike height, and the repolarization duration, which we defined as the time between the peak of the second spike and the local minimum of the respective afterhyperpolarization (see Methods for details). All responses were measured in the soma of the cell, both in experimental data and in our simulations.

To test our hypothesis — that both electrical and morphological properties contribute to neuronal response variability within the same cell type — required a multi-step approach. Firstly, we applied a large database of parameter sets to the same reconstructed morphology of a T cell to assess the contribution of variable electrical properties. Secondly, three example parameter sets were applied to a population of 15 reconstructed T cell morphologies that differed in morphological details, such as diameters and length of cellular processes (supplementary figure 1 and 2), to quantify the contribution of their morphological details to response feature variability. Since the spatial distribution of ion channels across the membrane of T cells is unknown, we compared different channel distribution configurations in the third step of the analysis to test whether our results depended critically on a specific location of the spike initiation zone. Finally, we investigated whether the experimentally observed systematic differences in response features between two subtypes (1RP and 2RP) of T cells could be explained by the morphological differences alone or if they require a difference in total membrane conductance.

**Figure 1.**
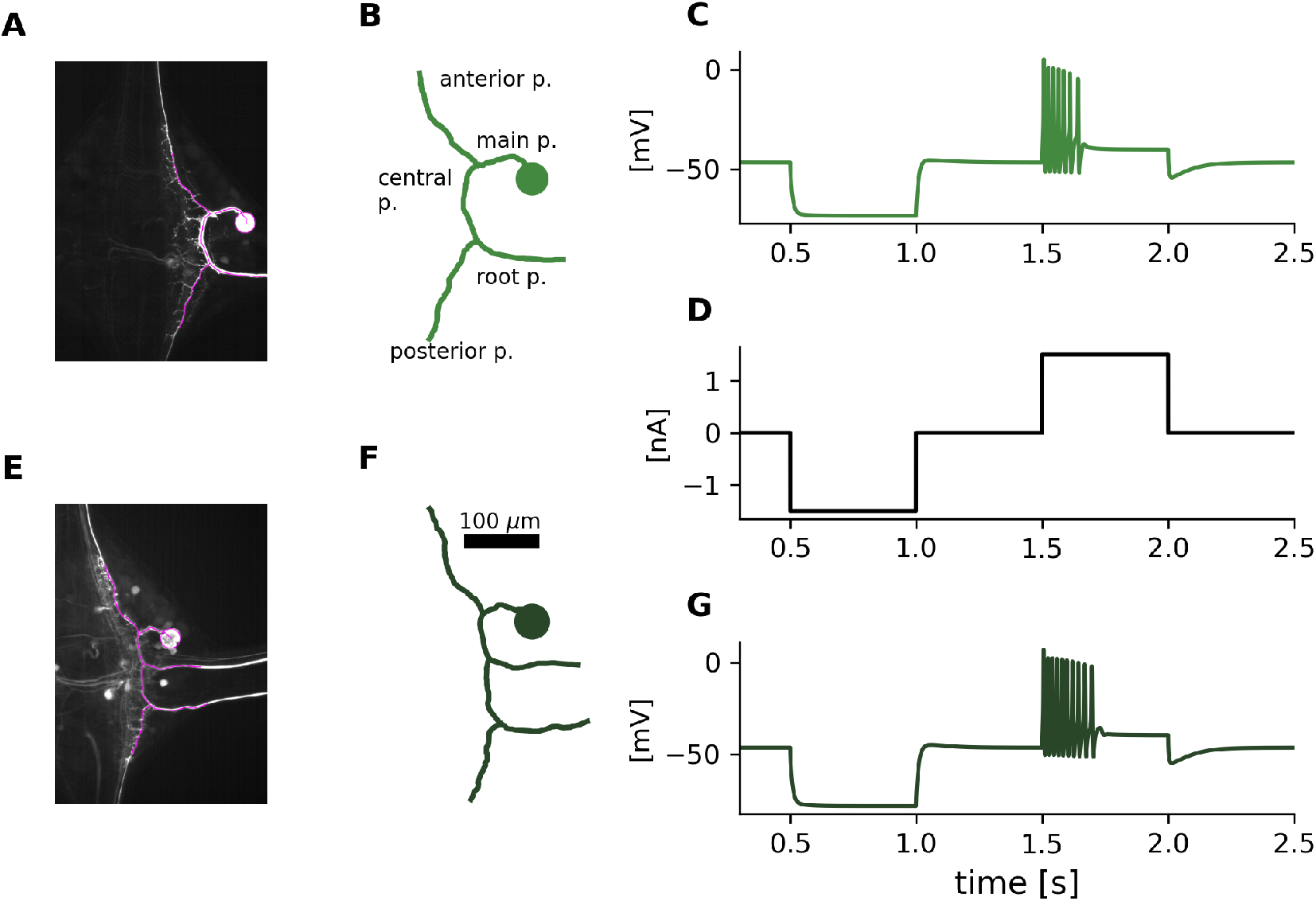
T cell morphologies and simulated responses. (**A**) Image of a 1RP T cell stained with neurobiotin. Magenta lines indicate the part of the morphology that was reconstructed. (**B**) Morphological reconstruction of the leech T cell shown in A), in which the SIZ was located in the root process, according to the standard configuration. Diameters of compartments are not drawn to scale here, see reconstruction 5 in supplementary figure 1 for a figure version indicating compartment diameters. The main branching pattern consisting of soma, main process, central process, anterior process, posterior process, and one root process, is the same for all 1RP morphologies (supplementary figure 1 shows all reconstructed morphologies with their total membrane area and total length). (**C**) Simulated response to somatic current injection of this reconstructed T cell morphology. The response was measured in the soma. (**D**) Stimulus protocol of the current injection (500 ms long pulses of -1.5 nA and 1.5 nA) into the soma, which was used in experiments and simulations throughout this study. (**E-G**) Cell image, reconstruction and simulated response for a 2RP morphology. See reconstruction 5 in supplementary figure 2 for a figure version indicating diameters. The main branching pattern of all 2RP morpholgies consist of the same processes as the 1 RP morphologies, plus an additional anterior root process (S2 Fig).

**Figure 2.**
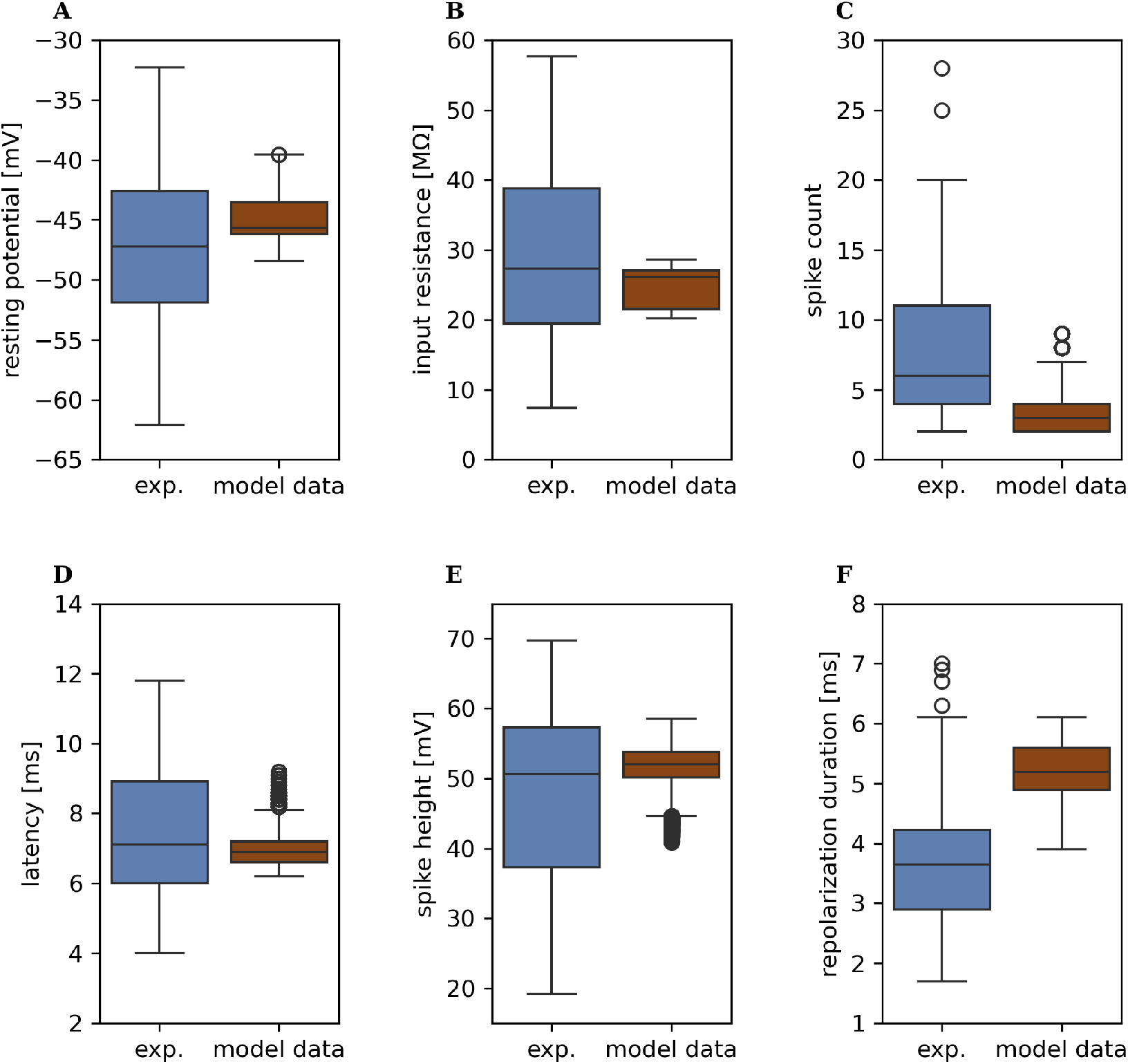
Response variability caused by electrical diversity applied to the same reconstructed morphology. Boxplots comparing response feature values of experimentally recorded n = 60 T cells (blue) to the model responses obtained for n = 33,958 parameter sets applied to the reconstructed morphology shown in Fig 1B (brown). The shown response features are (**A**) resting membrane potential (3 outliers in model data), (**B**) input resistance, (**C**) spike count (in 500 ms, 2 outliers in experimental data, 281 outliers in model data), (**D**) first spike latency (4 outliers in experimental data with very long latencies (27.7 ms, 20.3 ms, 16.5 ms and 15.1 ms)are not shown for improved visibility, 777 outliers in model data are shown), spike height (448 outliers in model data), (**F**) repolarization period (4 outliers in experimental data).

### 2.1 Many parameter sets reproduce T cell responses

We began the parameter sweep using a multi-compartment model of a 1RP T cell, with the SIZ placed in the single process which innervates the leech’s skin. Fig 1A shows the original microscopic image of this T cell. To ensure the same branching pattern and similar total length in all reconstructed morphologies, all fine branches were removed and the anterior, posterior and root process were trimmed (see Methods), reducing the microscopic image stack to the parts highlighted in magenta, and resulting in the reconstructed morphology displayed in Fig 1B. We tested 48,828,125 parameter sets (all combinations of 11 parameters with 5 values each, Table 2) for this morphology and found that 1,859,170 sets yielded valid responses. Fig 1C shows an example of a biologically realistic response simulation obtained with the reconstructed morphology shown in Fig 1B. We then applied these valid parameter sets to a population of 15 models based on individual reconstructed morphologies (see supplementary Figures 1 and 2), which all possess one of the two basic branching patterns shown in Fig 1 and were trimmed to similar length. 33,958 of the parameter sets generated valid responses for all reconstructed morphologies.

To isolate the contribution of electrical diversity on response variability, we applied all of these 33,958 parameter sets to the morphological reconstruction shown in Fig 1B. Different response features exhibited distinct degrees of variability (Fig 2). The variability of the resting membrane potential in the simulations obtained with this individual morphology covered 30% of the range observed in the experiments with n=60 T cells (Fig 2A). The range of input resistance represented 17% of the range observed in the experiments (Fig 2B). Our model generated spike counts ranging from 2 to 9 spikes for all valid sets applied to this reconstructed morphology (Fig 2C). Although approximately 75% of the experimentally observed data were found in this range, some of the experimental recordings yielded much higher spike counts, even up to 28 spikes in one cell. The simulated first spike latencies ranged from 6.2 ms to 9.2 ms for this reconstructed morphology, covering 13% of the variability observed in experiments (Fig 2D). The variability of spike height in the simulations represented 35% of the range from the smallest to the largest spikes in our datasets (Fig 2E). Simulated repolarization duration spanned from 3.9 ms to 6.1 ms, corresponding to 42% of the experimental range and covering rather the longer durations (Fig 2F).

Our results show that a biologically plausible T cell response can arise from many different electrical parameter sets in combination wit one specific morphological reconstruction. This supports the notion that T cells exhibit electrical diversity, contributing to the observed cell-to-cell variability in neuronal responses.

The variability in the simulated responses did not cover the entire experimentally observed range for any of the response features. This could be expected because the tested electrical parameter space was limited to five different values for each conductance and reversal potential parameter, and no ion-channel kinetics were varied. Hence, varying additional electrical parameters could introduce even higher response variability when applied to the same reconstructed morphology.

### 2.2 Morphological details contribute to response feature variability

Following the analysis variability introduced by multiple parameter sets applied to a single morphology, we next analyzed the variability caused by differences in morphological details. According to our criteria, 33,958 of the tested parameter sets generated plausible T cell responses for the entire population of 15 reconstructed morphologies. In this population, we included both morphological subtypes of T cells, eight 1RP and seven 2RP cells. The supplementary figures 1 and 2 provide an overview of all reconstructed morphologies. Applying the same parameter set to the population allowed us to isolate the impact of morphological details on response variability. Here, we illustrate this approach with three different parameter sets (Table 2 in Methods) which yielded systematically different spike counts, representing T cells with high, medium, and low activity, respectively (Fig 3).

**Figure 3.**
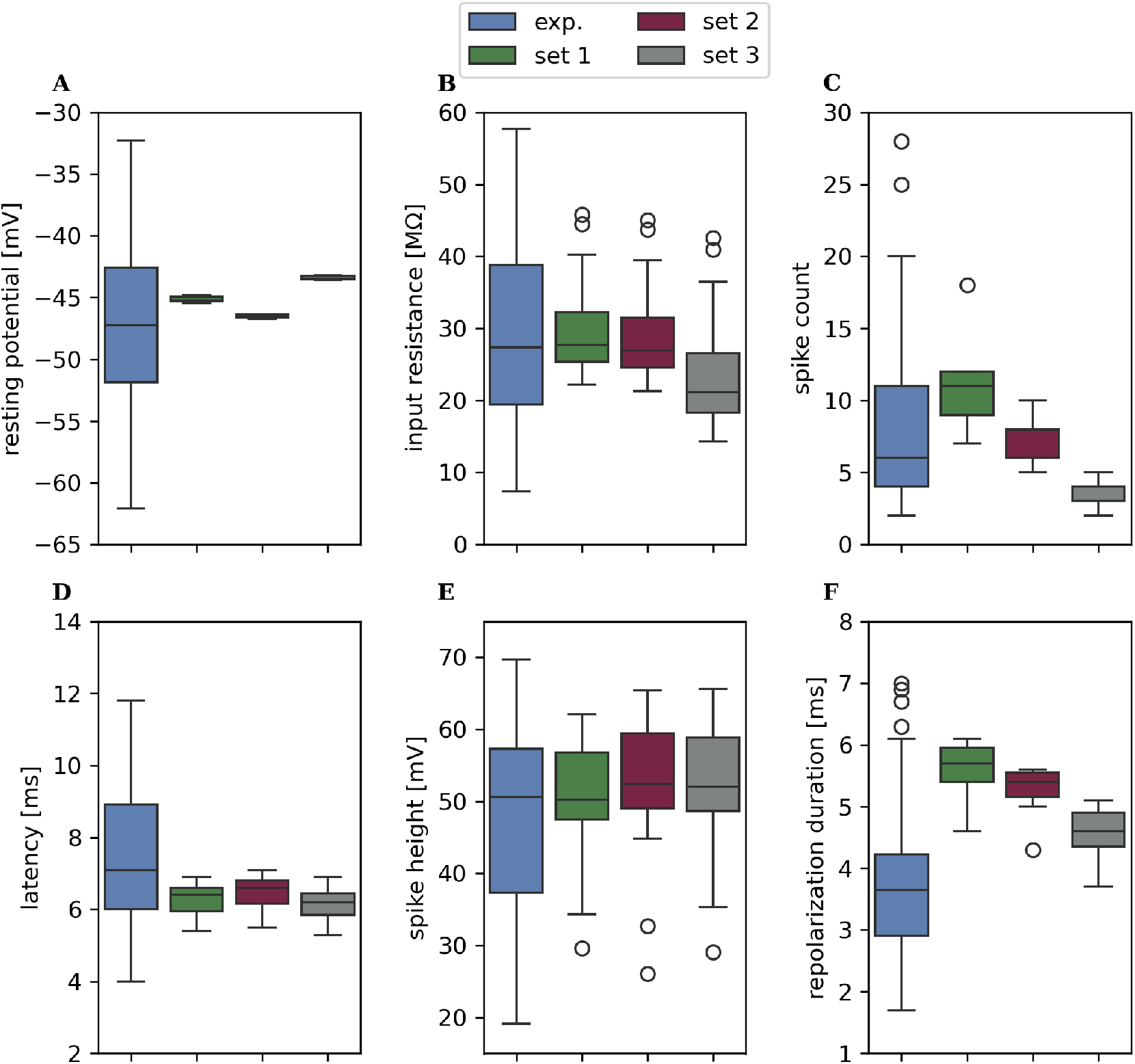
Variability in response features caused by morphological diversity with the same electrical properties. Boxplots of response features of experimentally recorded T cells (n = 60, blue, re-plotted from Fig 2) and three example parameter sets (green, red and grey) applied to n = 15 different reconstructed morphologies. The range of each boxplot shows the variability in response features introduced by morphological details of the reconstructed morphologies. The shown response features (with scaling as in Fig 2) are (**A**) resting membrane potential, (**B**) input resistance, (**C**) spike count (in 500 ms), (**D**) first spike latency (without outliers in experimental data, as in Fig 2D) (**E**) spike height, (**F**) repolarization duration.

The variability observed in the resting membrane potential was only ∼1% of the experimental finding for all three parameter sets, suggesting that cell morphology does not noticeably influence this response feature (Fig 3A). In contrast, the range of input resistance in each of our example sets represented ∼50% of the range observed in experiments, which indicates a high impact of morphological properties on the input resistance (Fig 3B). The range of spike count values was 11 for the parameter set representing highly active T cells (set 1, green), 5 for the parameter set yielding medium spike activity (set 2, red), and 3 for the parameter set causing low activity (set 3, grey) (Fig 3C). Each of the parameter sets covered only a fraction of the experimentally observed range, which stretched from 2 to 28 spikes in response to 500 ms of current injection. This suggests that morphological variety within a cell population contributes to variability of spike counts, but not as much as electrical differences do. The range of the simulated first spike latencies corresponded to only ∼ 7% of the range observed in experiments, and ∼20% of the range without outliers (Fig 3D, outliers are not shown). The height of the second spike was highly variable across reconstructed morphologies and covered ∼78% of the observed range in experimental data (Fig 3E). The repolarization duration was systematically above the experimentally observed median for all three parameter sets and covered on average 26% of the experimentally determined range for each of the parameter sets (Fig 3F).

We next investigated whether the observed variability in response features across individual reconstructed morphologies can be explained by a trivial effect. It is well known that the total membrane surface area of neuron impacts the neuron’s input resistance and thereby also its excitability (Kernell, 1966; Kernell and Zwaagstra, 1981). Therefore, all reconstructed morphologies in the population were truncated to the same distances between the soma and the tips of the anterior and posterior branches, respectively, in order to control for effects of cell size. Nevertheless, differences in the diameters of individual compartments and segment lengths resulted in a range of total surface areas and total lengths for both morphological subtypes (supplementary figure 1 and 2). Figure 4 illustrates how the response features depend on the total membrane surface area of the 15 reconstructed morphologies for parameter set 2. Remarkably, the total membrane surface areas (shown on the x-axis in all panels) of the two morphological subtypes did not fall into two distinct categories, but overlap substantially, although 2RP morphologies possess an additional root process compared to 1RP morphologies. Nevertheless, the total surface area of an individual 1RP reconstructed morphology with large compartment diameters can exceed that of a 2RP morphology with smaller diameters. While the resting potential was independent of the membrane surface area (Fig. 3A and Fig. 4A), the input resistance exhibited the expected strong negative linear dependence on the membrane − surface area (Fig. 4B, *r* = −0.91, *p* = 0.0002). However, this did not result in a correlation between membrane surface area and spike count, where the linear regression yielded a low *r*-value of *r* = 0.02, *p* = 0.47 (Fig. 4C). The spike response features of latency, spike height and repolarization duration were all positively correlated with the total membrane surface area, with a correlation factor between *r* = 0.68 and *r* = 0.79 (Fig. 4D-F). An increase in latency can be explained by a combination of the larger membrane capacitance, leading to slower voltage change, and the smaller input resistance leading to a smaller amplitude of the passive response to current injection. The smaller passive response also explains why the spike height is larger in cells with larger surface area. The greater the difference between the *Na*^+^ reversal potential and the membrane potential of the stimulated cell is, the larger the spikes can be. Repolarization duration, defined as the time difference between the peaks of spike depolarization and afterhyperpolarization, increases with spike height. Nevertheless, none of these considerations can explain why individual responses obtained with identical electrical parameters are distributed around the regression line. These differences are caused by the details of the reconstructed morphologies.

**Figure 4.**
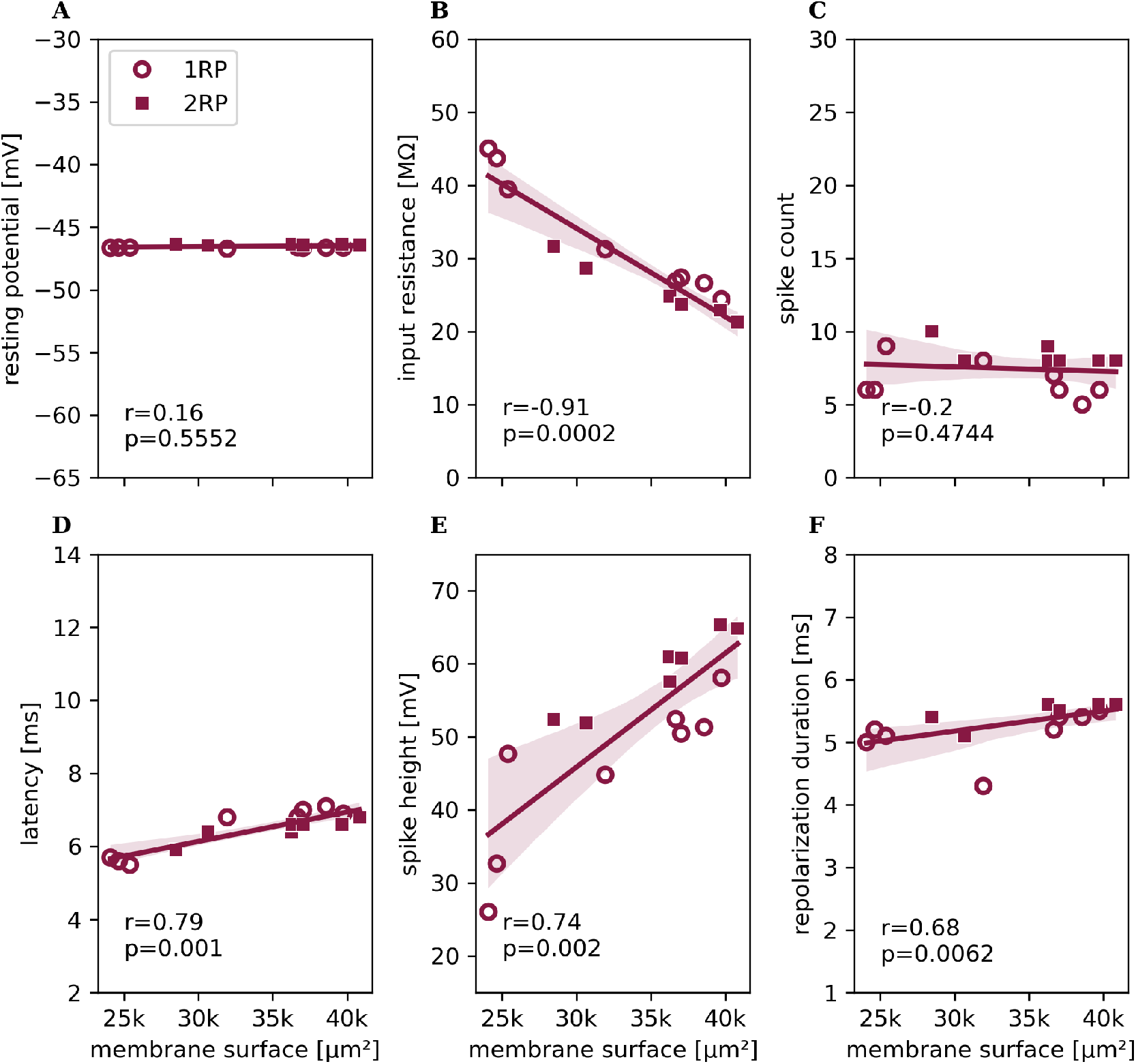
Dependence of response features on total membrane surface area. Response feature values obtained for all n = 15 different reconstructed morphologies with parameter set 2, plotted against the total membrane surface area of each reconstructed morphology. Feature values obtained for 1RP-morphologies are depicted with round symbols, 2-RP morphologies with square symbols. Lines show linear regression, shadows indicate the 95% confidence interval. For each response feature, *r* is the spearman correlation coefficient, and the *p* value is given below. The shown response features (with scaling as in Fig 2) are (**A**) resting membrane potential, (**B**) input resistance, (**C**) spike count (in 500 ms), (**D**) first spike latency, (**E**) spike height, (**F**) repolarization period.

To further investigate the interaction between spike generation and morphological details, we analyzed the rheobase of a population of 15 reconstructions to measure excitability. Our simulations reveal, that the rheobase of the reconstructions does significantly correlate with the membrane surface, but not with the input resistance (supplementary material 3).

In summary, even when the same parameter set was applied to the population of 15 reconstructed morphologies, morphological details alone introduced considerable variability to the responses of T cell models. These differences in response features could neither be explained by trivial effects caused by differences in the total membrane surface areas, nor by systematic differences between the two branching patterns (1RP and 2RP). Taken together, our results suggest that electrical diversity and morphological variation are both necessary to explain the observed range of response variance in T cells.

### 2.3 SIZ location influences spike features

As the location of spike generation in leech T cells is unknown, we tested whether our assumption of a SIZ location in the root branches was critical for the modeling results. This was analyzed separately for the two morphological subtypes. Starting with the reconstructed 1RP morphology presented in Fig 1A, we tested six different SIZ configurations (configurations i-vi in Fig 5A-F) using example parameter set 2. In the standard configuration (configuration i in Fig 5A, which was also used in all previous figures), the SIZ with larger conductance density was located in the root process. The other configurations included relocating the SIZ to one of the other processes of the cell (Fig 5B-E), as well as distributing the total conductance of the standard configuration homogeneously over the simulated membrane area (configuration vi in Fig 5F). All tested configurations generated spikes in response to somatic stimulation, although not all features remained within experimentally observed ranges. Compared to the standard configuration (i), spike counts can be higher or lower, depending on the SIZ location. The spike height measured in the soma differed as well between configurations. Spike amplitudes were larger when the SIZ was located in the main process (configuration ii, Fig 5B), or when the total conductance of the SIZ (Table 2) was distributed homogeneously across the entire cell membrane (vi) (Fig 5B and F).

**Figure 5.**
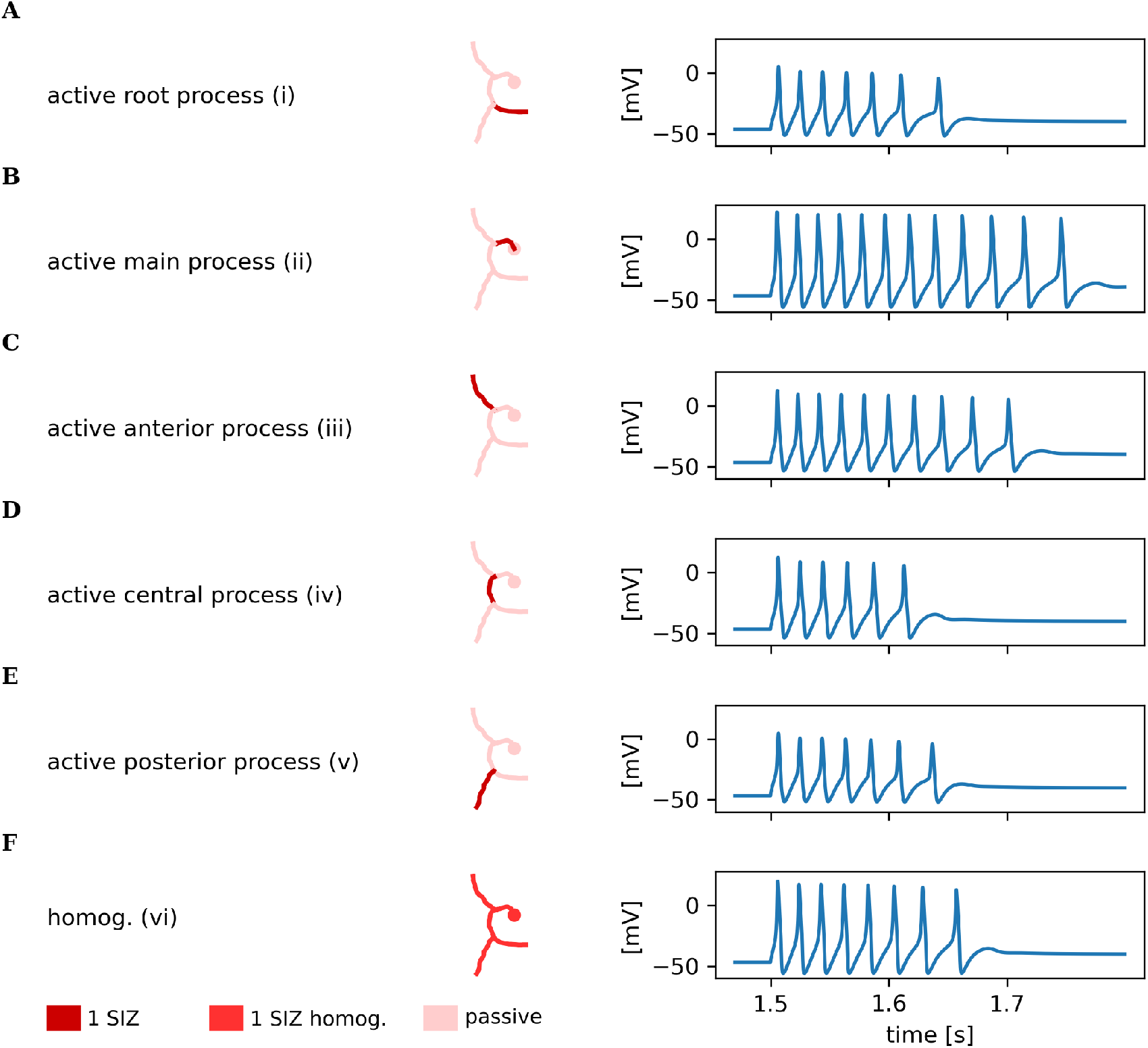
SIZ configuration in the same 1RP morphology influences spike responses. Example voltage traces measured in the soma compartment for six different SIZ configurations in the reconstructed 1RP morphology from Fig 1B. Conductance densities in the membrane of the reconstructed morphology are color coded with saturated colors indicating higher densities and areas of passive membrane shown less saturated (see colorbar). (**A**) SIZ located in the root process to the skin is the standard 1RP SIZ configuration used in this study. (**B**) SIZ located in the main process (configuration ii, close to the soma) results in more and larger spikes in the soma. (**C**) SIZ located in the process to the anterior ganglion (configuration iii) increases spike height and spike count slightly compared to standard configuration i. (**D**) SIZ located in the central process (configuration iv) leads to a similar number of spikes with slightly increased height compared to configuration i. (**E**) SIZ located in the process to the posterior ganglion (configuration v) elicits a similar spike count and spike height as the standard SIZ configuration i. (**F**) Distributing the conductances of the SIZ section homogeneously over the whole membrane (configuration vi) causes larger spikes.

When analyzing the impact of the SIZ location on response features for all reconstructed morphologies of our population, we found that the SIZ location has virtually no impact on the resting membrane potential or the input resistance of the simulated responses (not shown). For the spike features, the influence of the SIZ configuration was variable across the reconstructed morphologies. Figure 6 color-codes the differences between the response feature values of each SIZ configuration (ii-vi: x-axes) and the standard configuration i for each reconstructed 1RP morphology in the population sorted by surface area (y-axes corresponding to numbers in supplementary figure 1). The spike count increased in all reconstructed morphologies when the SIZ was located either in the main process (ii) or in the anterior process (iii), with morphologies with smaller total membrane surface showing the strongest effects (Fig 6A). The strongest increase in spike count was observed for the reconstructed morphology 1 (the morphology with the smallest total surface area) in configuration ii, which responded with 24 additional spikes. When the SIZ was located in the central- or posterior process (configurations iv and v), only minor changes in the number of spikes were observed. When the total conductance was distributed homogeneously (vi), most reconstructed morphologies responded with more spikes compared to their standard configuration, except for the reconstructed morphology 4, which produced one spike less.This reconstructed morphology differs from the other members of the population in having a thick and long central process, while the other processes are rather thin (see supplementary figure 1). The first spike latency decreased strongly for configurations ii and iii, in which the SIZ was located closer to the soma, as well as the homogeneous conductance distribution vi (Fig 6B). The latency of reconstructed morphology 1 reduced by 2.4 ms when the SIZ was located in the main process (configuration ii), making it faster than any experimentally observed T cell. As with spike count, configurations iv and v had only minor effects, producing slight decreases, no change, or, for the morphology with the largest surface, a negligible increase in latency. Spike height increased for all 1RP models when the SIZ was located in the main process (ii), in the anterior process (iii), or in the central process (iv) (Fig 6C). Reconstructed morphology 1 showed an increase of spike height by 48.23 mV for configuration ii, exceeding the experimentally observed spike height in our dataset. Placing the SIZ in the posterior process (v) yielded similar spike heights compared to the standard configuration. When conductance densities were distributed homogeneously (configuration vi), the spike height increased for all morphologies, similar to configuration ii. The impact of the SIZ location on the repolarization period was variable across reconstructions, but 7 out of 8 reconstructions showed shorter repolarization periods when the SIZ was located in the distal process (v), or when conductance densities were distributed homogeneously (vi). The highest increase of 0.9 ms was observed in reconstructed morphology 4 (with a large central process and thin other processes) in configuration iii and the largest decrease of -1 ms for morphology 8 (with the largest surface) in the homogeneous configuration vi.

**Figure 6.**
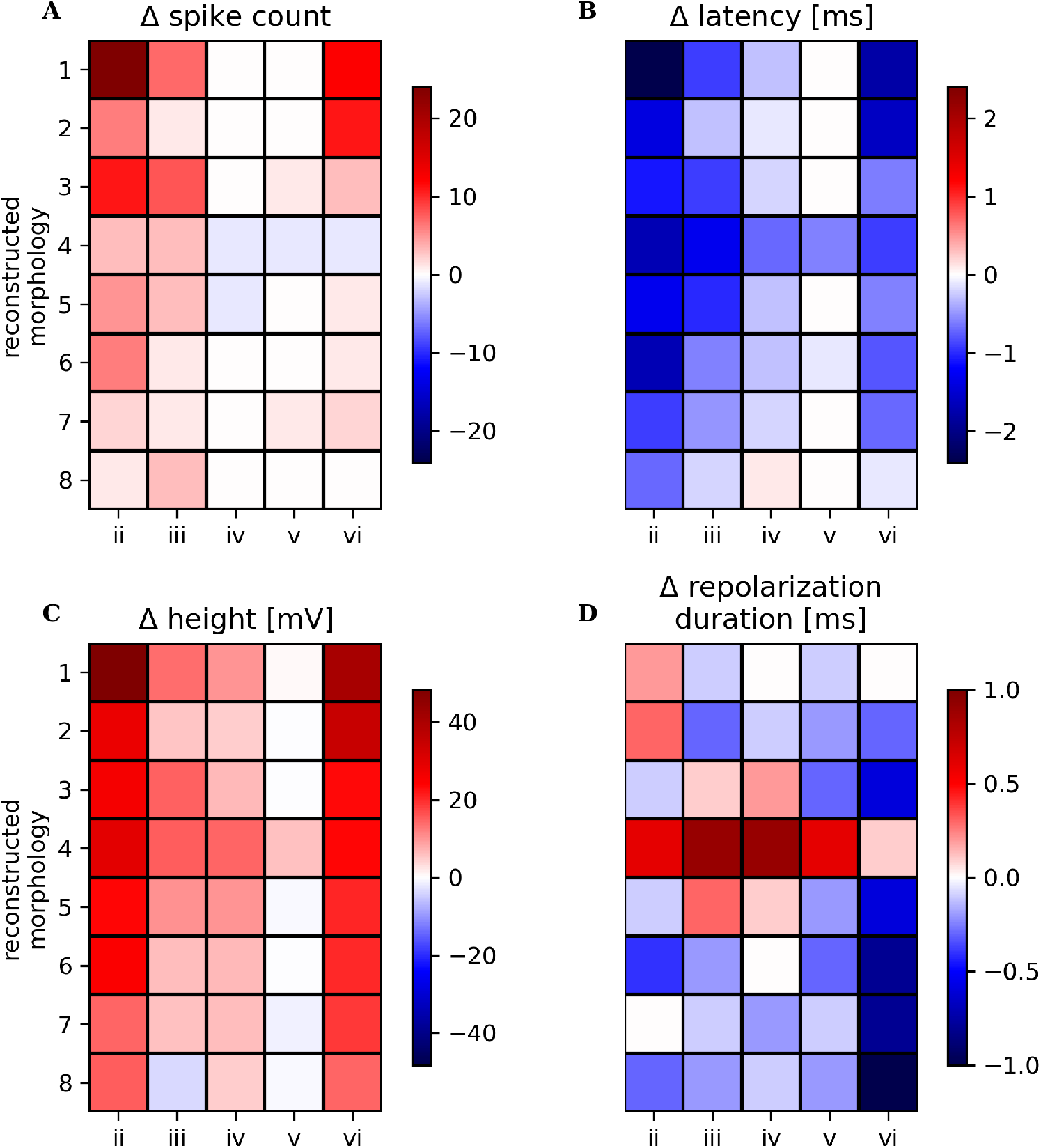
SIZ configuration influences spike response features in all reconstructed 1RP morphologies. Heatmaps of spike feature differences depending on the SIZ configuration (x-axes) for 8 reconstructed 1RP morphologies sorted from smallest surface area (1) to largest surface area (8) (y-axes), morphology 5 is the same as shown in Fig 5. Labels of SIZ configurations refer to those in Fig 5. Differences were calculated for each reconstructed morphology by subtracting the response feature value obtained for the standard SIZ configuration i (Fig 5A) from the value obtained for each SIZ configuration. Differences in spike counts, (**B**) first spike latency, (**C**) spike height, and (**D**) repolarization duration are color coded with increases shown in red and decreases in blue (see colorbar).

For the population of seven 2RP T cells we assumed in the standard configuration (I in Fig 6A, also used in Figs 1 and 3) that they have two SIZs, one in each root process. This assumption is based on the findings of Meiser et al. (Meiser et al., 2023), that 2RP cells have systematically higher spike counts and spike heights compared to 1RP cells, which lead to their hypothesis that 2RP cells have two SIZs, while 1 RP cells have only one SIZ. We tested six SIZ configurations for 2RP models with example parameter set 2 and one example reconstructed morphology (configuration I-VI in Fig 7A-F, same morphology as in Fig 1 and number 5 in supplementary figure 2) to investigate how the distribution of active membrane affects the response features of a 2RP T cell. In configuration II, we placed one SIZ in the anterior process and one in the posterior process, to separate them spatially as far as possible from each other, without changing the total conductance (Fig 7B). In this configuration, we observed higher spikes compared to the standard configuration I.

**Figure 7.**
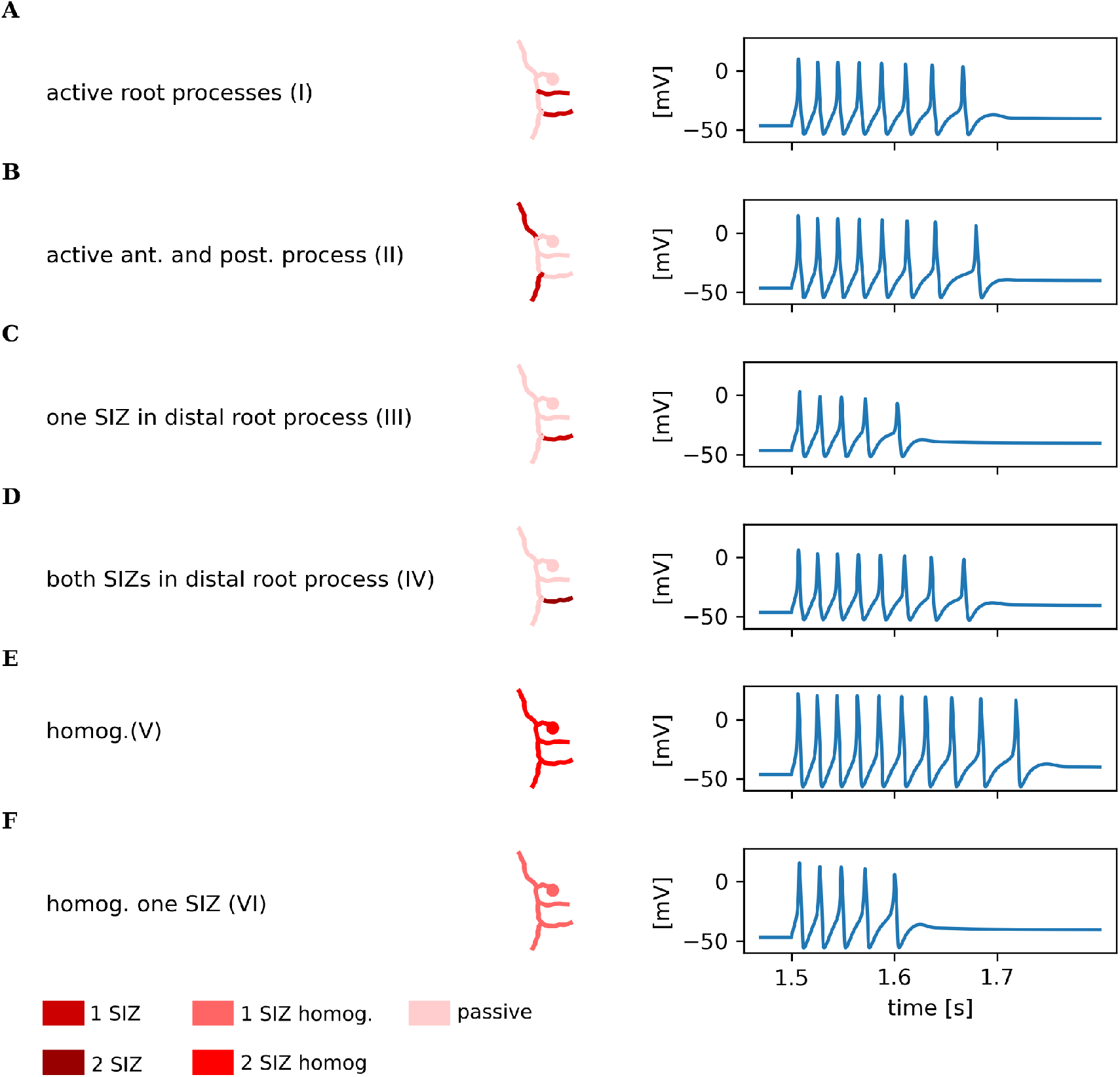
SIZ configuration in the same 2RP morphology influences spike responses. Voltage traces resulting from different SIZ configurations in an example reconstructed 2RP morphology, displayed as in Fig 5. (**A**) Standard 2RP configuration: SIZs located in both processes to the skin. (**B**) SIZs located in the anterior and posterior processes elicit a similar spike count and slightly higher spikes. (**C**) Omitting the SIZ in the proximal process to the skin and keeping the SIZ in the other root process as in the standard configuration results in a reduced spike counts and smaller spikes compared to the standard configuration for 2RP (A), and also for 1RP (Fig 5A). (**D**) Combining the conductance of both SIZ in the distal root process elicits a similar response to the standard configuration. (**E**) Total conductance of configuration IV homogeneously distributed increases the spike count and spike height. (**F**) Total conductance of configuration IV (approximately half of the total conductance compared to V) homogeneously distributed elicits fewer but higher spikes than the standard configuration I.

With configuration III, we challenged our assumption of 2RP T cells having two SIZs. In this configuration, we omitted the SIZ from the proximal root process. This left the reconstructed morphology with approximately half the total conductance of standard configuration I for 2RP morphologies, and a similar amount to 1RP morphologies, which by definition have only one SIZ. This reduction of total conductances caused fewer and smaller spikes in the simulated responses for configuration III (Fig 7C) compared to the 2RP morphology in configuration I, and even compared to the standard configuration of the 1RP morphology (compare Fig 5A). When the total conductances from both SIZ in the standard configuration were combined in one SIZ in the distal root process (configuration IV), the simulated response was similar to the standard configuration, demonstrating that the total conductance, rather than the number of SIZs, is the primary factor of excitability (Fig 7D).

To investigate the impact of total conductances separately from morphological properties, we considered two configurations in which the total conductance of SIZs was distributed homogeneously across the membrane of the reconstructed morphology. In configuration V, we distributed the total conductance of the standard configuration I homogeneously over the cell membrane and observed more and larger spikes in the simulated response, compared to the standard configuration I (Fig 7E).

Lastly, we homogeneously distributed the total conductance of configuration III (only one SIZ) across the reconstructed morphology (configuration VI in Fig 7F), resulting in a homogeneous configuration with approximately half the total conductances compared to configuration V. For this homogeneous distribution of a lower total conductance (configuration VI), we observed fewer, but larger spikes compared to the responses of the standard configuration I and considerably fewer spikes than for configuration V with a homogeneous distribution of higher total conductance.

Analogue to the analysis of 1RP reconstructions, we calculated the difference between the response feature values of each SIZ configuration and the standard configuration I for each 2RP reconstructed morphology (Fig 8). Spike counts decreased in all reconstructed morphologies for the two configurations with lower total conductances, i.e. when the SIZ in the proximal root process was omitted to allow a direct comparison to 1RP reconstructions (III) or when the conductance density of configuration III was distributed homogeneously in configuration VI (Fig 8A). When the conductance of the standard configuration I was distributed homogeneously (V), spike counts increased for all reconstructed morphologies, with a stronger impact on morphologies with smaller total surface. Placing SIZs in the anterior and posterior process (II) caused a slight increase of spike counts in two reconstructed morphologies and a slight decrease in two others. When the conductance densities of both SIZs were combined in the distal root process (IV), responses were similar to the standard configuration I. First spike latency decreased in most reconstructed morphologies, when SIZs were located in the anterior and posterior process (II), and also when SIZ conductances of both SIZs were distributed homogeneously in configuration V (Fig 8B). Omitting one SIZ (III), or distributing the conductance of one SIZ homogeneously (VI) increased the first spike latency. Combining both SIZs in the distal root process (IV) led to similar first spike latencies compared to the standard configuration. The observed spike height decreased, when the SIZ in the proximal root process was omitted (III). This effect remained, when the conductance of both SIZ were combined in the distal root process in configuration IV (Fig 8C). Remarkably, the spike height increased compared to the standard configuration when the conductances of only one SIZ were distributed homogeneously (VI), although the total conductance was lower. Repolarization period became shorter for all configurations, except for configuration II where both SIZ were shifted to compartments that were further apart (Fig 8D).

**Figure 8.**
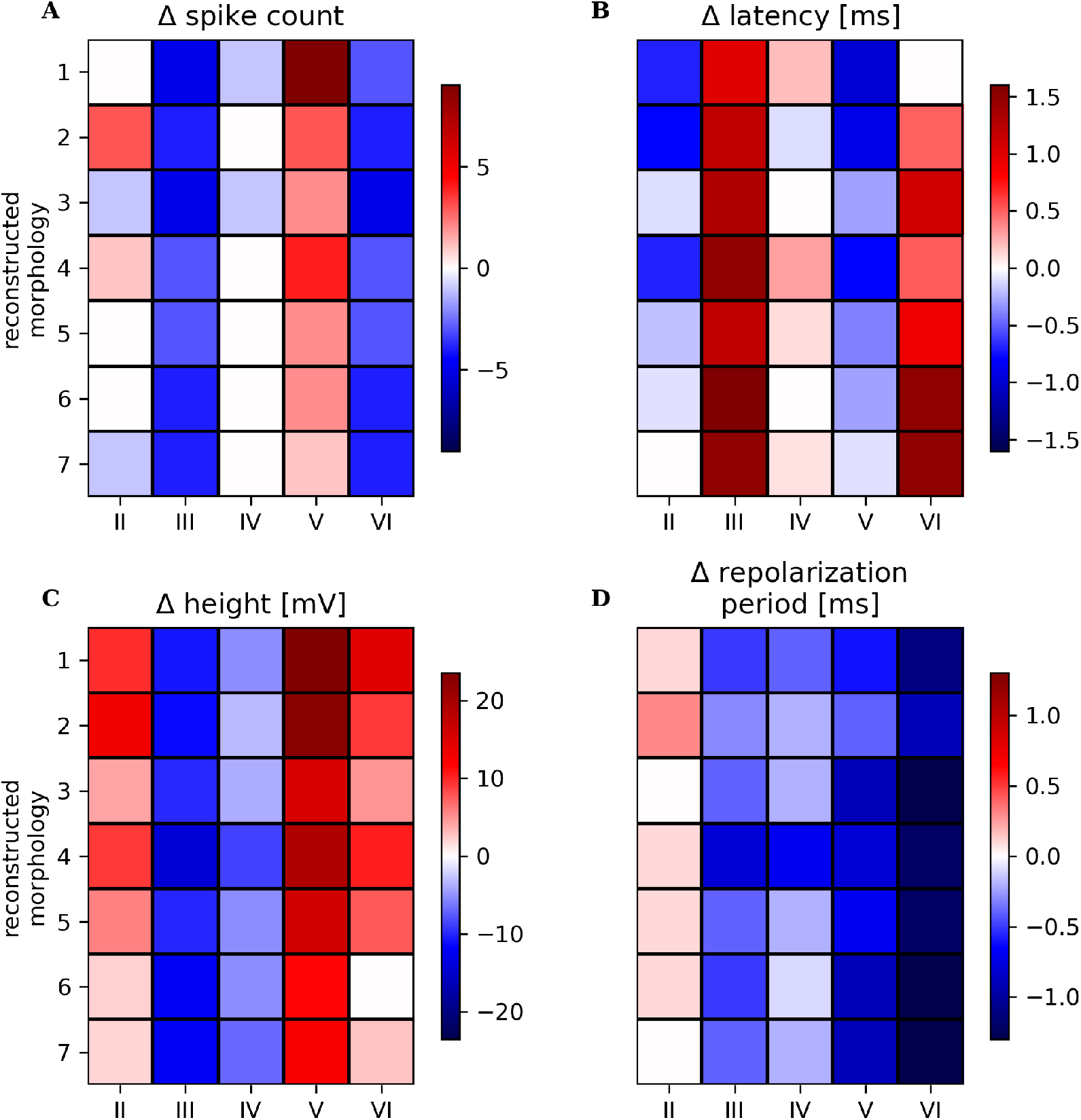
SIZ configuration influences spike response features in all reconstructed 2RP morphologies. Heatmaps of differences in response feature values compared to the 2RP standard configuration I (Fig 7A) were calculated accordingly as in Fig 6 for 7 reconstructed morphologies with two root processes and sorted from smallest surface area (1) to largest surface area (7), morphology 5 is the same as shown in Fig 7. Labels of configurations on the y-axis refer to those in Figure 7. Differences in (**A**) spike counts, (**B**) first spike latency, (**C**) spike height, and (**D**) repolarization duration are color coded with increases shown in red and decreases in blue.

By testing different SIZ configurations for both morphological subtypes we found that they did not impact the passive response features (input resistance and resting potential), but strongly influenced all spike features. It should be kept in mind that the parameter sweep was performed based on the assumption of the standard SIZ configurations and therefore combined eight 1RP reconstructed morphology with one SIZ and seven 2RP reconstructed morphologies with two SIZ. Our results show for one example parameter set that biologically plausible responses do not critically depend on this experimentally not testable assumption, but can qualitatively also be obtained with other SIZ configurations, although all spike response feature values depend quantitatively on the SIZ configuration (Fig 6 and 8).

### 2.4 Systematical differences between morphological subtypes can be explained by total conductance of active membrane

In our exploration of the parameter space, we assumed one SIZ for T cells with 1RP morphology and two SIZs for the 2RP morphological subtype as standard configurations. Therefore, it is important to determine whether differences in morphology or in total conductance can explain the higher activity of the 2RP morphological subtype observed in experiments.

In the strip plots of Figure 9, the experimental data set from Figure 2 is compared to the simulation results obtained for the three parameter sets of Figure 3 applied to the eight 1RP and the seven 2RP reconstructed morphologies. All three example parameter sets showed the same trends: The 2RP morphologies in the standard configuration I generate more and higher spikes than the 1RP morphologies in standard configuration i when the same parameter set is applied (compare same colors in Fig 9A). This result is consistent with the experimental findings by Meiser et al. (Meiser et al., 2023), but depends critically on the SIZ configurations. For most neuronal types, it is assumed that they have only one SIZ, usually located at their axon initial segment (Huang and Rasband, 2018). When simulating all leech T cells, independent of their number of root processes, with only one SIZ with the same conductance density, (comparison of configurations i and III), the spike counts of the 2RP models fall below the spike counts of 1RP models with the same parameter set. Combining conductance densities from both SIZs into the distal root process (configuration IV), 2RP models respond with spike counts similar to their standard configuration I. These findings imply that it is very likely for T cells with two root processes to possess a higher total conductance than the morphological subtype with one root process to explain the experimentally found higher number of spikes in the 2RP morphological subtype of T cells.

**Figure 9.**
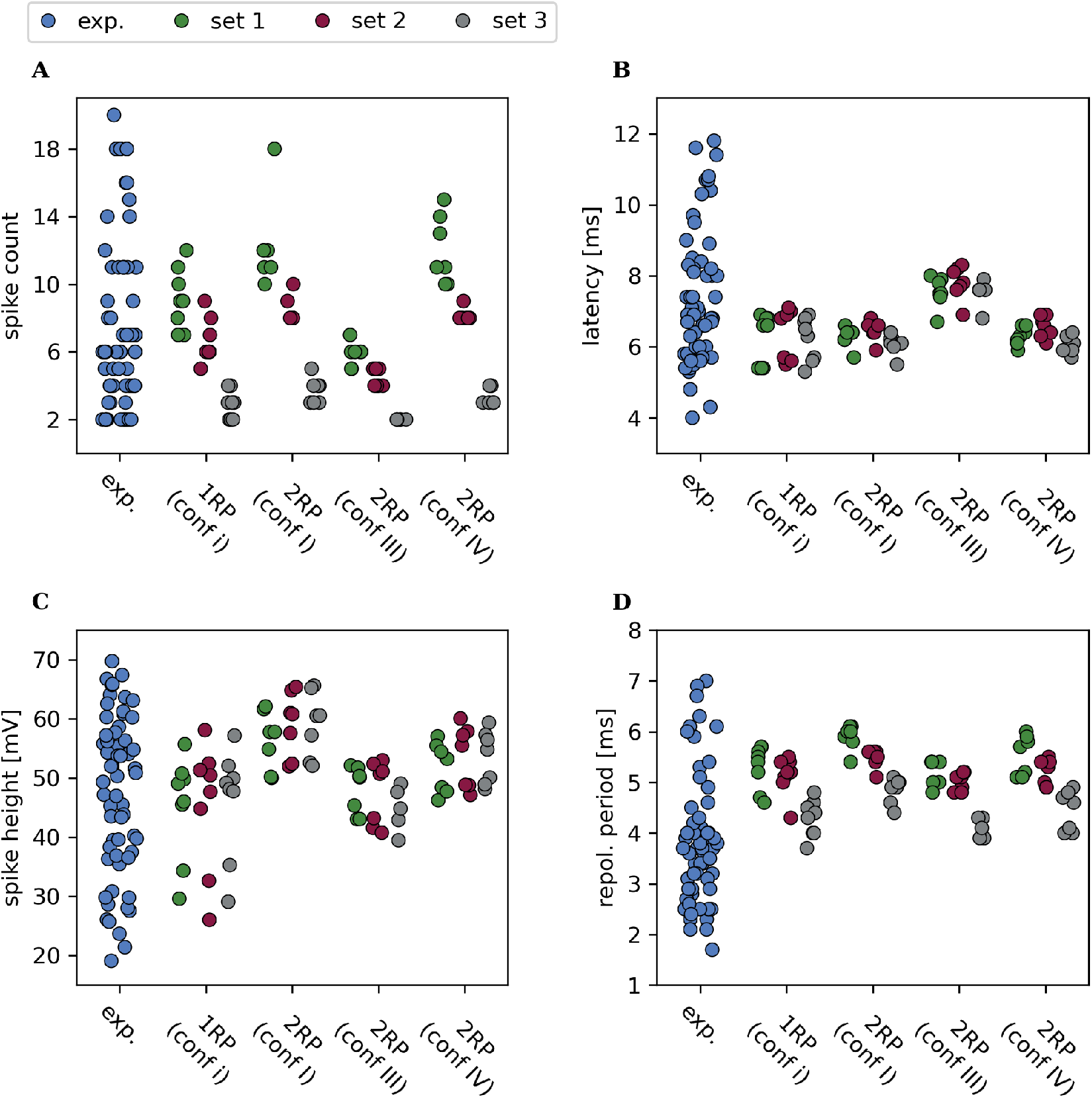
Systematic differences in spike responses between reconstructed 1RP and 2RP morphologies. Strip plots of spike response features of n = 60 experimentally recorded neurons (blue, same data as in Fig 2 and 3), and three example parameter sets (green, red and grey, same parameter sets as in Fig 3) applied to n = 8 one root process morphologies with an SIZ located in their root process (see Fig 5A, standard configuration i, and supplementary figure 1 for reconstructed morphologies) and n = 7 two root process morphologies with two SIZs in each root process (see Fig 7A, standard configuration I, and supplementary figure 2 for reconstructed morphologies). Additionally, response features for the 2RP morphologies are shown for SIZ configurations III (one SIZ with half total conductance, Fig 7C) and IV (one SIZ with full total conductance, Fig 7D). For each SIZ configuration, the color indicates the applied parameter set. Strip plots for (**A**) spike counts, (**B**) first spike latency, (**C**) spike height and (**D**) repolarization period.

In an experimental study (Meiser et al., 2023), first spike latencies were reported to be shorter for the 2RP morphological subtype. In our small samples of reconstructed morphologies, we did not observe a consistent latency difference between 1RP and 2RP models under their standard SIZ configurations (Fig 9B). However, when the SIZ in the proximal root process was omitted in configuration III, the latency increased beyond the values observed for the 1RP models, which would contradict the experimental findings. This latency increase was due to the lower total conductance, as can be seen from the comparison with configuration IV, in which the standard total conductance condensed into one SIZ leads to approximately the same latencies as observed for the standard configuration I (Fig 9B).

Spike height was systematically larger for 2RP morphologies in configurations I and IV compared to 1RP morphologies (Fig 9C). When the proximal SIZ was omitted, which reduced the total conductance (configuration III), 2RP spike heights decreased to the 1RP level. In configuration IV, condensing the total conductance of configuration I into one SIZ increased spike height relative to configuration III, but values remained below those of the standard 2RP configuration I. These findings also indicate that also the experimentally found differences in spike height (Meiser et al., 2023) depend strongly on the total conductance. The repolarization periods, which were not investigated in the experimental study (Meiser et al., 2023), followed the trends observed for spike heights and spike counts (Fig 9D).

These results indicate that the previously reported differences in spike counts and spike heights between 1RP and 2RP T cells cannot be explained by morphology alone, but instead require differences in total conductance and its distribution. Assuming approximately the same total conductance for both morphological subtypes leads to results that contradict the experimental findings. While our model study clearly shows the importance of a higher total conductance in the 2RP morphological subtype, it is still unanswered if the total conductance is distributed to two separate SIZ located within the central ganglion, since other SIZ configurations lead to similar results.

## 3 DISCUSSION

Understanding the sources of variability in neuronal responses remains a fundamental challenge in neuroscience. This study demonstrated that variability in morphological details, even within the same cell type and fixed bran pattern, does impact neuronal response features. Using a large database of parameter sets that generated biologically plausible leech T cell responses across 15 reconstructed morphologies (Fig 2), we demonstrated that differences in morphological details like diameters and length of cellular sections induce variability in T cell responses (Fig 3). The observed range of response features, in particular spike counts, cannot be explained trivially by differences in total membrane surface area between the reconstructed morphologies (Fig 4 and supplementary figure 3). Our results indicate that previously reported typical responses of 1RP and 2RP T cells (Meiser et al., 2023), can arise from different SIZ configurations (Fig 5 and 7) and that the distribution of ion channels in the neuronal membrane has an impact on spike features (Fig 6 and 8). Our approach also offers an explanation for the systematic differences which may be driven by differences in SIZ properties between the two morphological subtypes of 1RP and 2RP (Fig 9).

### 3.1 The interplay of morphological details, electrical diversity and channel distribution as sources of variability

Our study focused on the influence of electrical and morphological diversity on two subthreshold response features and four spike response features of leech T cells. Changes in SIZ configuration had virtually no effect on input resistance, whereas varying electrical properties or introducing morphological diversity produced considerable variability. Resting membrane potentials in the simulations were less variable than in experiments, suggesting that they are only weakly influenced by electrical or morphological variability or by SIZ configuration. However, it should be noted that the range of experimentally observed resting membrane potentials in this data set is larger than in previous studies on T cells (Meiser et al., 2019; Scherer et al., 2022). It is likely that the process of impaling the cell membrane during the intracellular recording impacts the resting membrane potential during experiments, and therefore the experimental data may overestimate the variability of the resting membrane potential compared to unperturbed T cells. Additionally, the variability in simulated resting potentials is likely underestimated, since the membrane capacitance *C*_*m*_ and the axial resistance *R*_*i*_ were not varied in this study.

Spike features showed different dependencies on the three factors of variability, electrical diversity, differences in morphological details and location of the SIZ. Electrical diversity caused variable spike counts, as reported in previous studies Günay et al. (2008); Berger and Crook (2015), but did not make the example reconstructed morphology in Fig 2 a very active T cell. However, high spike counts were found for other morphologies in the population of simulated T cells (Fig 3C). While differences in membrane surface and input resistance partly explain the observed variability in latency, spike height and repolarisation period (see Fig 4B and D–F), they do not completely account for the differences in excitability, nor do they explain the differences in spike counts between individual morphologies or morphological subtypes (see Fig 4C and Fig 5). This confirms that morphological details are an important factor in neuronal excitability. The location of the SIZ also influenced the spike count strongly, suggesting that the exact location of the SIZ might be variable from T cell to T cell. First spike latency showed little variability between different electrical parameter sets, or between reconstructed morphologies. The SIZ configuration, however, strongly affected the first spike latency, especially in reconstructions with a smaller surface area (Fig 6B and Fig 8B). This is in line with findings of a previous modeling study, reporting that spike latency of vertebrate neurons is mostly influenced by cell size and SIZ location (Gulledge and Bravo, 2016). Spike timing carries a lot of information in mechanosensation (Johansson and Birznieks, 2004), and therefore our results underline the crucial role of SIZ location in T cells, and may also apply in other mechanoreceptors. Spike height was highly variable when one parameter set was applied to the population of reconstructed morphologies and when the SIZ location was varied, while it was less strongly impacted by the electrical parameters. We found variable repolarization durations for all three factors we investigated, but the repolarization duration was systematically above the median value of experimentally observed responses. For homogeneously distributed ion channel conductances, we found shorter repolarization periods in almost all morphological reconstructions. This could indicate that we underestimated the ion-channel conductances in the passive compartments. Another possible factor that could influence the repolarization period as well as the other spike features is the ion channel kinetics (Kole et al., 2007; Goaillard and Marder, 2021b; Schneider et al., 2023), which were not varied in this study, but only approximated based on a patch clamp recording (supplementary figure 4).

Our results agree with the findings of prior studies on degeneracy in neural systems that many distinct parameter combinations can produce functionally equivalent outputs (Prinz et al., 2004; Achard and De Schutter, 2006; Alonso and Marder, 2019). We identified 33,958 parameter sets that generated responses resembling T cells for 15 reconstructed morphologies. Morphological properties such as the soma size and location, dendritic arborization and axonal properties are known to shape neural response features (Brette, 2013; Hesse and Schreiber, 2015; Fékété et al., 2021). Our results reveal that morphological details alone can give rise to variability in response features such as input resistance, spike count, and spike height, even when all electrical parameters and the overall branching pattern are held constant. The finding that the total membrane surface area correlates only weakly with most response features and is uncorrelated with spike counts suggest that even minor morphological differences, such as the diameter and length of a dendritic branch, contribute to the response variability of T cells in a non-trivial way. These complex effects of morphological details probably affect neuronal response variability in general, and may therefore contribute to variability at the network level (Gowers and Schreiber, 2024). Moreover, our approach likely underestimated the impact of morphology, since we used simplified reconstructions of morphologies and therefore did not take the fine arborization into consideration, which is also a known source of variability (Cuntz et al., 2010; Peng et al., 2021).

It is clear that there are more sources of neuronal response variability than morphological details, the electrical parameters of conductances, and the density and spatial distribution of ion-channels, which we considered in this study. Other factors including the ion channel dynamics (Goaillard and Marder, 2021b), chemical synaptic (Roffman et al., 2012) and electrical coupling between cells (Crodelle et al., 2019) are also known to contribute to neuronal variability. Further studies should determine how all of these mechanisms interact to give rise to robust networks and facilitate many possible solutions for functional output.

### 3.2 Systematic differences between subtypes of a defined cell type

Understanding how morphological properties influence neuronal responses is a crucial task in computational neuroscience, especially for classifying and understanding the function of subtypes of neuronal cell types (Chand et al., 2015; Kole and Brette, 2018; Peng et al., 2021; Scala et al., 2021; Moubarak et al., 2022; Mao and Staiger, 2024).

Leech T cells are a well-studied and experimentally accessible example of neurons with two morphological subtypes. All T cells are mechanoreceptors, which respond to light touch on the skin. Previous experimental and modeling studies suggested that all T cells require at least two SIZs to reliably transmit signals from the skin to the central part of the ganglion: a peripheral SIZ is located in the skin, and a central SIZ is located close to the ganglion (Burgin and Szczupak, 2003; Kretzberg et al., 2007). Here, we simulated a T cell in an isolated ganglion, neglecting the distal processes that innervate the skin with the peripheral SIZ. Unfortunately, experimental limitations do not allow us to measure ion channel distribution on the membrane of T cells and therefore prevent the exact SIZ location from being determined. The electrophysiological recordings allow only indirect reasoning: Since the first spike in response to current injection into the soma occurs at a median latency of 7 ms, the SIZ is likely not close to the soma. This argument is supported by a previous model study, in which a one-compartment model failed to reproduce the T cell response latency (Meiser et al., 2019). Our simulation result for the 1RP morphological subtype points in the same direction. When the SIZ was located directly next to the soma in the main process, the simulated latencies were shorter than those observed in experiments (Fig 6B). Nevertheless, since several different configurations of channel distributions, including SIZ locations in different segments, as well as homogeneously distributed ion channels, reproduce biologically plausible responses, our approach was unable to determine the location of the SIZ in biological T cells. Our results indicate that the location of the central SIZ is important for spike timing and other spike response features, although it may vary from cell to cell. It is also possible that the location of the SIZ in T cells might change dynamically and thereby contribute to homeostatic plasticity, a mechanism proposed by prior studies to regulate the excitability of cells in response to sensory or synaptic input (Grubb et al., 2011; Yamada and Kuba, 2016; Goldstein et al., 2019).

In this study, we compared two morphological subtypes, which are known to differ also in function. In each ganglion, two T cells of the 1RP morphological subtype innervate the dorsal region of the leech skin, while the other four T cells possess a 2RP morphology and respond to touch on the ventral or lateral skin regions (Nicholls and Baylor, 1968). The distribution of ion channels across the membrane is even more unclear for the 2RP than for the 1RP morphological subtype. Since both root processes reach the skin at different positions, it is safe to assume that both of them possess a peripheral SIZ that send action potentials towards the ganglion when the respective skin area is stimulated. It is unknown if this morphological subtype also posses two central SIZ. But our simulations show that the experimental observation that T cells with two root processes exhibited higher spike counts and greater spike amplitudes than their one-root counterparts (Meiser et al., 2023) can only be explained by a higher total conductance in the 2RP morphological subtype. If the same total conductance is assumed in both subtypes, the larger membrane area of the additional root process reduces the excitability of 2RP models, so that they produce fewer instead of more spikes than the 1RP models. Hence, the 2RP morphological subtype must possess a greater number of ion channels, but our model approach does not allow a conclusion if these are located in one densely populated SIZ, split between two SIZs, or distributed across a larger membrane area.

Through the manipulation of SIZ properties and locations, we showed that increased active membrane area and proximity to the soma can enhance excitability. This finding suggests an interplay between morphology and electrical properties that explains subtype-specific functional differences. This supports the hypothesis that additional SIZs in the root processes could contribute to the systematic differences between responses of T cell subtypes. Our results reveal how morphology, ion channel distribution, and electrical properties interact to generate variability in neuronal responses, highlighting multiple pathways by which the nervous system achieves reliable function.

## 4 MATERIALS AND METHODS

We wrote custom software in Python 3.9.13 for data analysis and simulations. T cell models were designed using Brian 2, version 2.5.4 (Stimberg et al., 2019).

### 4.1 Data analysis of electrophysiological data

Data from **N=60** cells of the previously published T cell dataset from Meiser et al. (Meiser et al., 2023) were reanalyzed. The test protocol was 11 s long and consisted of a -1.5 nA pulse for 500 ms starting at 4 s and a 1.5 nA pulse for 500 ms starting at 8.5 s (Fig 1C). We only considered the second spike of the response for spike shape features, because the shape of the first spike differs systematically from subsequent spikes. Therefore, we excluded all cell responses with a spike count of 1. We analyzed the following response features:

- **Resting membrane potential** [mV] was determined as the median membrane potential over 500 ms before the first stimulation.
- **Input resistance** [MΩ] was calculated as follows:

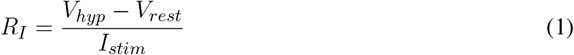
- with *R*_*I*_ as the input resistance, *V*_*hyp*_ as the median membrane potential during a 500 ms hyperpolarizing current pulse of *I*_*stim*_ = −1.5 nA and *V*_*rest*_ beeing the resting membrane potential.
- **Spike count** was defined as the number of action potentials in response to a 500 ms depolarizing current injection of +1.5 nA.
- **Latency** was determined as the time difference between the stimulus onset of the +1.5 nA current injection and the peak of the first spike.
- **Spike height** was measuered as the absolute voltage difference between the peak of the second spike in response to current injection of +1.5 nA and the minimum of the respective afterhyperpolarization within a time window of 10 ms.
- **Repolarization duration** was defined as the time difference between the peak of the second spike in response to +1.5 nA current injection and the minimum of the respective afterhyperpolarization.

The rheobase was chosen as a measurement of excitability (Rotterman et al., 2021). We applied incrementally increasing current pulse amplitudes from 0.1 nA and 1 nA in steps of 0.1 nA for 500 ms, and determined the rheobase as the first amplitude that elicited a spike. Since our experimental stimulus protocol was not designed for a rheobase measurement, we only determined the rheobase for simulated responses.

### 4.2 Morphological reconstruction

We used 15 of the anatomical reconstructions previously published by Meiser et al. (Meiser et al., 2023). Eight were reconstructions of T cells with one root process, and 7 were T cells with two root processes. The comparison between the microscopy pictures and the corresponding reconstructed morphologies is shown for two examples in Fig 1. The supplementary figures 1 and 2 display all 15 reconstructed morphologies, and indicate the diameters of their compartments. We applied a Ramer-Douglas-Peucker algorithm to all reconstructions to simplify structures equally and reduce variability in the number of compartments between cells (Hirschmann, 2014). We set the distance tolerance *ϵ* to 1 µm in the algorithm for all morphologies, resulting in a median of 120 compartments for one-root-process reconstructions and a median of 162 compartments for two root process models. To reduce methodological variability of morphologies, the minimum distance from the tip of the anterior process to the soma were determined and the anterior process of all morphologies were cut to match this value for all morphologies (see magenta lines in Fig1A and E). The same procedure was applied to the posterior process. Root processes were also cut to the shortest length found in the population to ensure same length in all morphologies.

### 4.3 Multi-compartment model

We reproduced electrical T-cell responses with a conductance based multi-compartmental model. The dynamics of the membrane voltage in each compartment was calculated as follows:

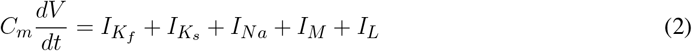

With *C*_*m*_ = 1 *µF/cm*^2^ as the standard membrane capacitance, *V* is the membrane voltage at time point *t* and *I*_*x*_ describes the transmembrane current for each current type. Compartments were connected by an intracellular resistance of *R*_*i*_ = 100 Ω *cm*, as in previous studies (Goethals and Brette, 2020). Voltage clamp experiments of our lab revealed fast and slow components in the *K*^+^ current (supplementary figure 4), so we added corresponding ion-channels in our model. Due to the large size of leech cells, only a rough estimation of the ion-channel kinetics was possible, which was used as a starting point for hand tuning of ion channel kinetics. For the fast *K*^+^ current, we implemented a Shaw-like Kv3.1 channel which is often expressed in cells that generate rapid trains of action potentials (Kanemasa et al., 1995). Channels of this type were also identified in the transcriptome of *H. verbana* (Northcutt et al., 2018). *We formalized the current of the fast potassium channel as follows:*

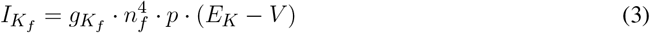

With 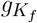 as the conductance density of the fast potassium channel, *n*_*f*_ and *p* are the gating variables, *E*_*K*_ is the reversal potential of K^+^ and *V* is the membrane potential. For the slow potassium current we defined:

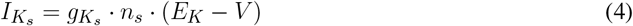

With 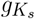 as the conductance density for the slow potassium channel and *n*_*s*_ is the gating variable. The fast transient sodium channel (Johansen, 1991) was formulated as:

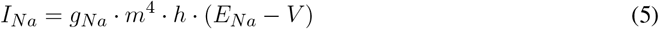

where *g*_*Na*_ is the conductance density for Na^+^-ions, *m* and *h* are the gating variables, *E*_*Na*_ is the reversal potential of Na^+^ and V is the membrane potential. An M-type potassium channel was added to cease spiking during current injection (Benda and Herz, 2003; Meiser et al., 2019). It was formulated as:

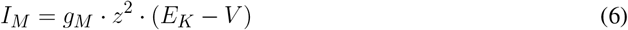

With *g*_*m*_ as the maximum conductance of the M-type channel and *z* as the gating variable. Kinetic equations for the slow and fast potassium channel where formulated as:

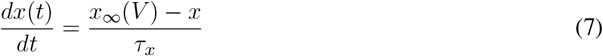

Where *x* is either *n*_*f*_, *n*_*s*_, *p, m, h*, or *z, x*_∞_ corresponds to the respective steady state value and *τ*_*x*_ is the respective time constant (Table 1). The steady state functions where defined by:

**Table 1.** Constant parameter values of ion-channel kinetics.

|  |  |  |  |
| --- | --- | --- | --- |
| $I_{K_f}$ | $n_{f1}; n_{f2} [mV]$ | 1; 16.5 | 8, 9 |
| $I_{K_f}$ | $p_1; p_2 [mV]$ | 15; 8 | 8, 9 |
| $I_{K_f}$ | $T_{n_f} [ms]$ | 1 | 9 |
| $I_{K_s}$ | $n_{s1}; n_{s2} [mV]$ | 25; 29.19 | 8, 9 |
| $I_{K_s}$ | $T_{n_s} [ms]$ | 67.18 | 9 |
| $I_M$ | $z_1; z_2 [mV]$ | 40.76; 2.62 | 8, 9 |
| $I_M$ | $T_z [ms]$ | 94.03 | 9 |
| $I_{Na}$ | $m_1; m_2 [mV]$ | 20; 11 | 8, 9 |
| $I_{Na}$ | $h_1; h_2 [mV]$ | 28; 10.35 | 8, 9 |
| $I_{Na}$ | $T_m [ms]$ | 0.8 | 9 |
| $I_{Na}$ | $T_h [ms]$ | 19.25 | 9 |

**Table 2.** Ranges of varied parameters. We chose 5 values for each varied parameter, resulting in 5^11^ = 48, 828, 125 combinations.

| Parameter | min | max | step | set 1 | set 2 | set 3 |
| --- | --- | --- | --- | --- | --- | --- |
| $E_K [mV]$ | -80 | -40 | 10 | -50 | -60 | -50 |
| $E_{Na} [mV]$ | 20 | 60 | 10 | 40 | 30 | 60 |
| $E_L [mV]$ | -60 | 0 | 15 | -45 | -45 | -30 |
| $g_{K_f} [mS/cm^2]$ | 250 | 1250 | 250 | 500 | 750 | 500 |
| $g_{K_f}(SIZ) [mS/cm^2]$ | 600 | 1140 | 180 | 600 | 960 | 960 |
| $g_{K_s} [mS/cm^2]$ | 0.01 | 0.09 | 0.02 | 0.09 | 0.03 | 0.07 |
| $g_{Na} [mS/cm^2]$ | 7 | 11 | 1 | 7 | 8 | 7 |
| $g_{Na}(SIZ) [mS/cm^2]$ | 140 | 380 | 60 | 140 | 200 | 140 |
| $g_M [mS/cm^2]$ | 1 | 5 | 1 | 1 | 1 | 3 |
| $g_M(SIZ) [mS/cm^2]$ | 0.1 | 1.7 | 0.4 | 0.1 | 0.5 | 0.1 |
| $g_L [mS/cm^2]$ | 0.02 | 0.1 | 0.02 | 0.1 | 0.1 | 0.08 |

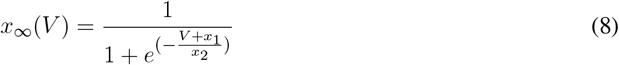

For the sodium and M-type potassium current, *τ*_*x*_ was voltage dependent and was calculated as:

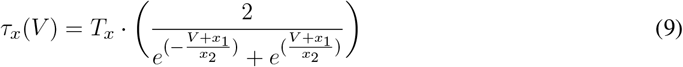

## 4.4 Spike initiation zone configuration

The SIZ had higher conductance densities for 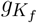, *g*_*M*_ and *g*_*Na*_ (see model section), compared to the other sections. In the standard configuration, the SIZ was located in the root process of the 1RP morphologies. For the 2RP morphologies, both root processes contained a SIZ with the same conductance density. Hence, the total conductance was higher in the standard configuration of 2RP than of 1 RP morphologies. For the homogeneous configurations vi and V (Fig 5F and Fig 7E) we calculated the total conductance of 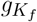, *g*_*M*_ and *g*_*Na*_ in the standard configuration and distributed it homogeneously over the membrane surface. For the homogeneous configuration VI (Fig 7F), we used configuration III and distributed its total conductance homogeneously.

### 4.5 Parameter study

We simulated responses to a shortened stimulation protocol for somatic current injection. We applied the same stimulus as described above for experimental data. To evaluate the response for a given parameter set, we compared each response feature to the corresponding experimentally derived data. A simulated response was considered valid if the following inequation was true for all analyzed response features:

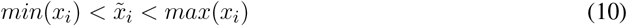

with *max*(*x*_*i*_) and *min*(*x*_*i*_) as the maximum respectively the minimum of the experimentally measuered response feature values and 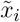 as the corresponding response feature value in the simulated response.

Using the Python package psweep, we created a parameter grid with all possible combinations of the given parameters (Table 2). The sweeps were parallelized on the HPC cluster of the University of Oldenburg, utilizing the Dask Python package (Dask Development Team, 2016). The parameter sweeps were done in two steps: **1**: All combinations of the parameters in Table 2 were applied to one T-cell morphology with one branch. The simulated responses were compared to the experimentally observed responses concerning the response features listed above. **2**: The parameter combinations that yielded valid responses for the selected example anatomy were tested on the entire population of morphologies. In this parameter search, a parameter set was considered valid if inequation 10 was true for all morphologies. For the examples presented in section 2.1, we decided to show a morphology with response feature values at the center of the response feature distribution for the entire population. For the example simulations in the “Results” section, we randomly selected three parameter sets that yielded different spike counts, mimicking highly, medium and less active T cells in our data set (Table 2).

## AUTHOR CONTRIBUTIONS

KS and JK: conceptualization, interpretation, and drafting manuscript. KS: Modelling and data analysis. IA: Anatomical sudies. JK: project administration and supervision. All authors contributed to the article and figures design, and approved the submitted version.

## ACKNOWLEDGMENTS

We thank the division for computational neuroscience and especially Dr. Go Ashida, Daniela Antonia Schwarz, Austin McNamara and Brittni Paige Devlin for valuable discussion and feedback on the manuscript.

## DATA AVAILABILITY STATEMENT

The datasets for this study can be found in the gin-gnode repository: [https://gin.g-node.org/Strandpalme/Sandbote_et_al_2026].

