## supplementary figures for "Morphological details contribute to neuronal response variability within the same cell type"

### Supplementary Material

#### 1 SUPPLEMENTARY FIGURES

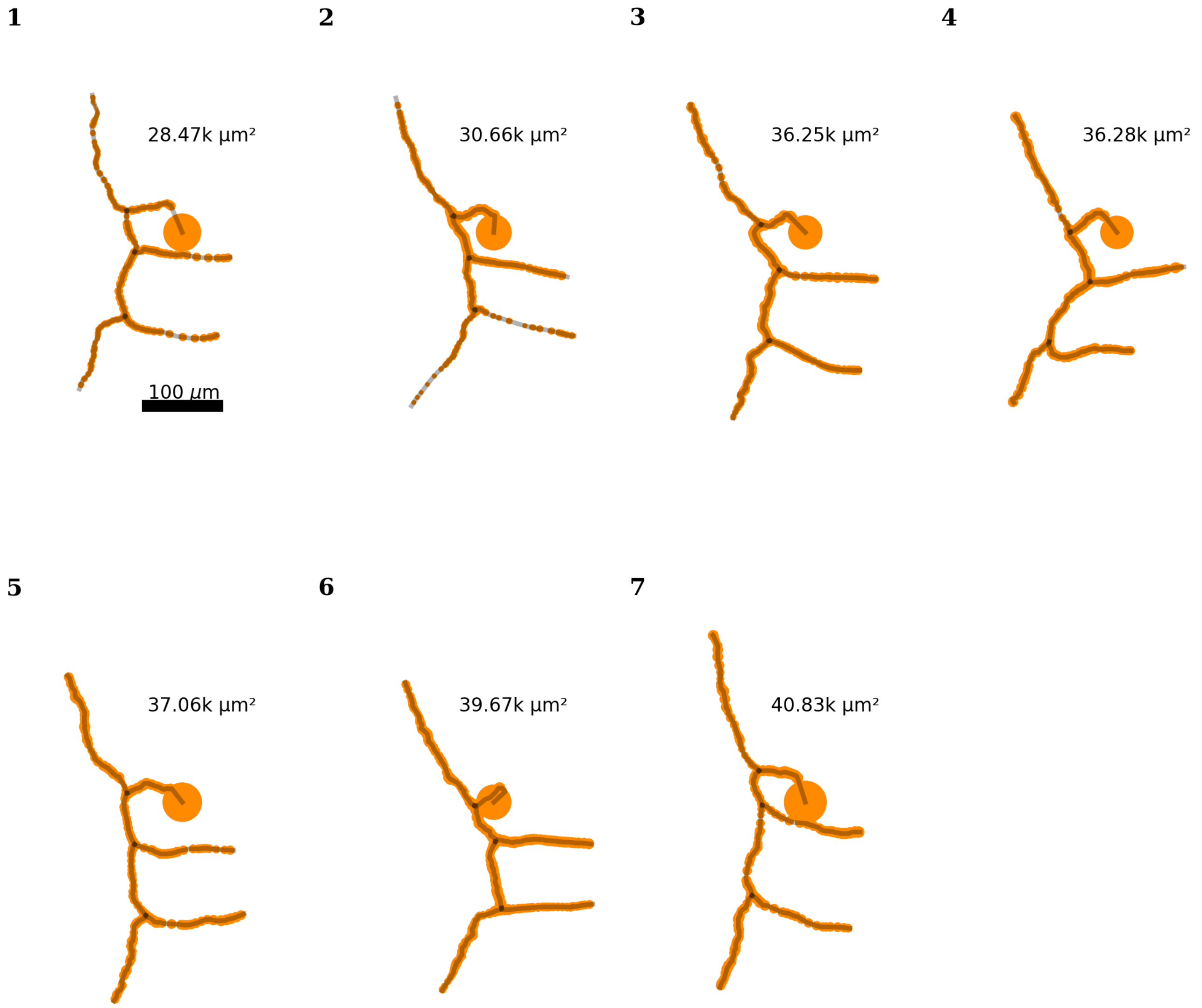


**Figure S1. Reconstructed 1RP morphologies.** All reconstructed 1RP morphologies used in this study, sorted by total membrane surface area (given in $\mu m^2$ in the panels) to match labeling in Fig 6. Gray lines indicate the skeleton of the reconstruction, the orange dots indicate the center and diameter of individual compartments.

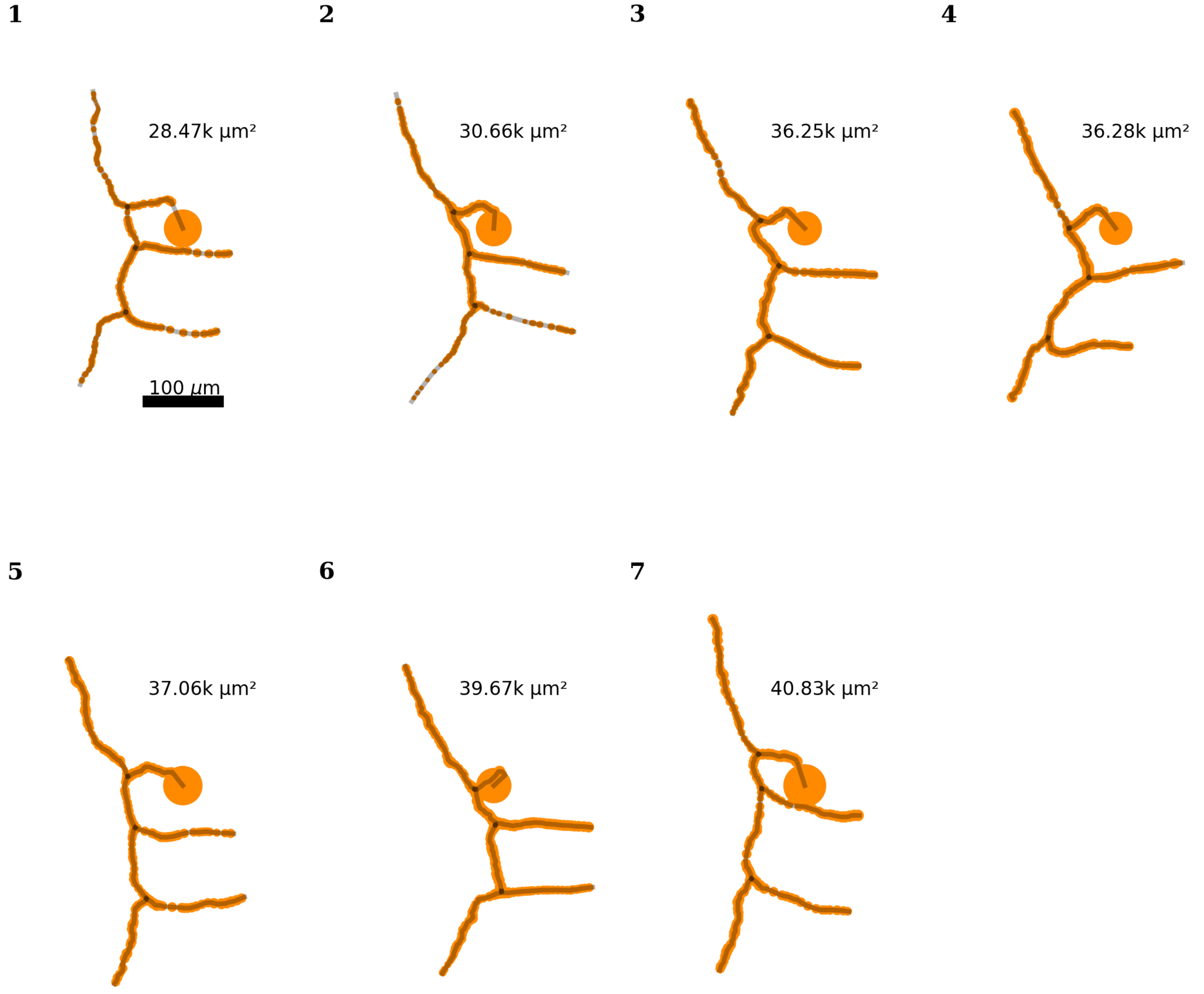


**Figure S2. Reconstructed 2RP morphologies.** All reconstructed 2RP morphologies used in this study, sorted by total membrane surface area (given in $\mu m^2$ in the panels) to match labeling in Fig 8. Gray lines indicate the skeleton of the reconstruction, the orange dots indicate the center and diameter of individual compartments.

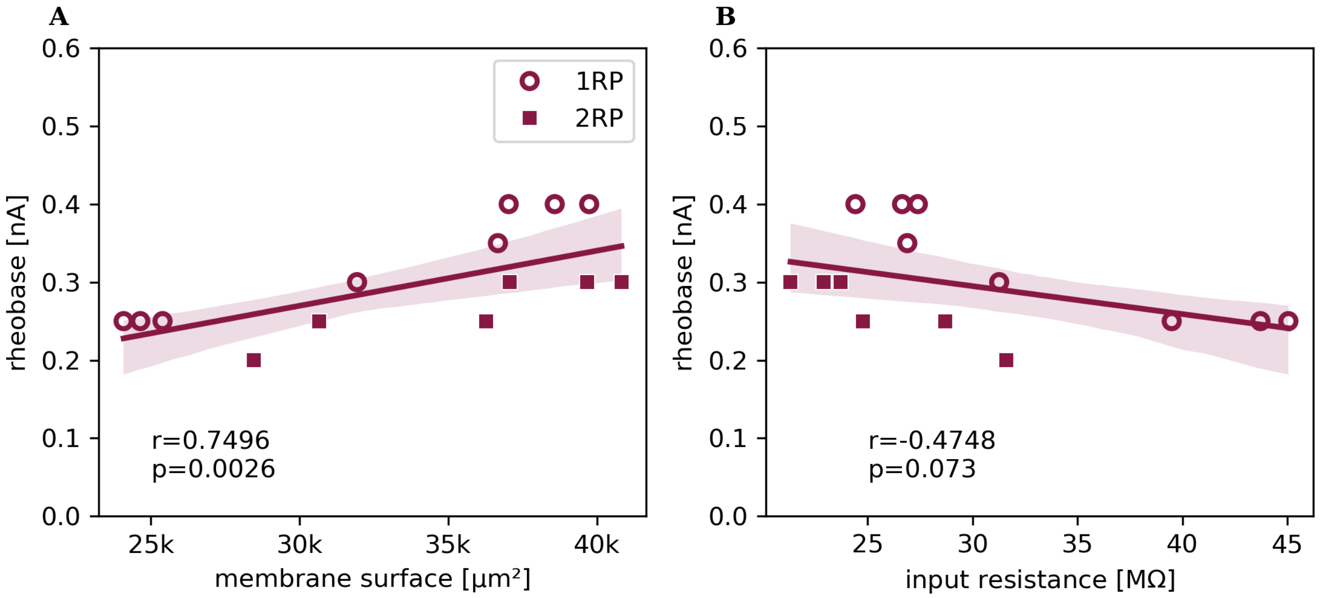


**Figure S3. Underlying factors of excitability.** The rheobase, defined as the minimum injected current that triggered a spike response, obtained for all n = 15 different reconstructed morphologies with parameter set 2, varies across morphologies. Like in Fig. **??**, rheobase values obtained for 1RP-morphologies are depicted with round symbols, 2-RP morphologies with square symbols. Lines show linear regression, shadows indicate the 95% confidence interval. For each response feature, $r$ is the linear correlation coefficient, and the $p$ value is given below. The rheobase is plotted against A) the total membrane surface area and B) the input resistance.

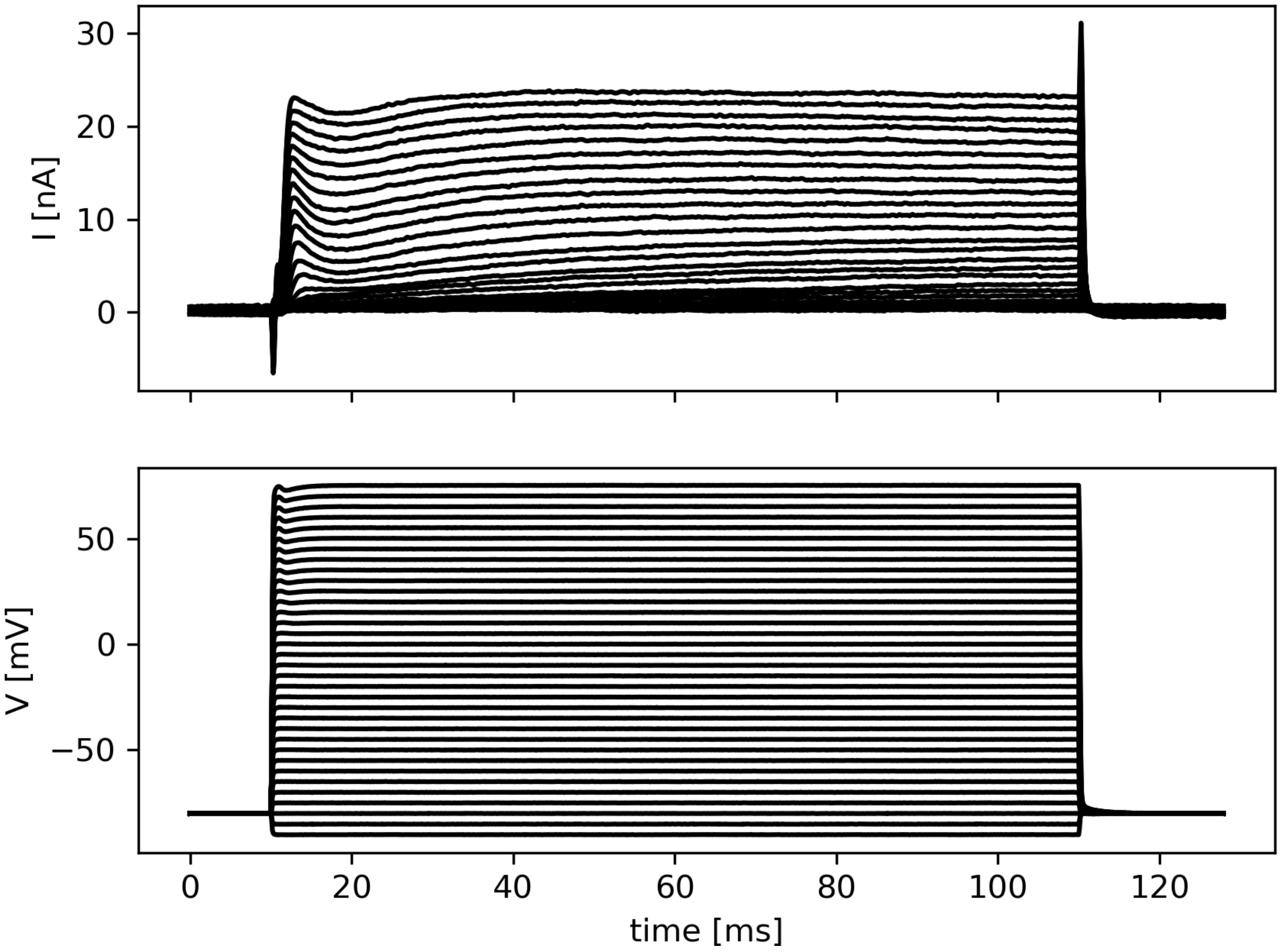


**Figure S4. Patch-clamp recording of potassium currents in a leech Touch cell in sodium-free solution.** Potassium currents were isolated by using a sodium-free solution. The top panel shows outward $K^+$ currents, each trace corresponding to a voltage-step protocol shown in the bottom panel. The cell was held at –90 mV, and depolarizing steps were applied in 5 mV increments up to +75 mV. At low voltages, the $K^+$ current exhibits a slow component that dominates throughout the 100 ms stimulation. With increasing depolarization, a fast-activating component emerges at the onset of the voltage step. Due to the large size of the T cell, complete space clamp could not be achieved, which limits the voltage control to the soma and proximal neurite. Therefore, the recordings represent an approximate measure of somatic and primary neurite potassium currents.
